# *In vivo* screening of TCR-based chimeric antigen receptors for improved anti-tumor functionality

**DOI:** 10.1101/2025.07.23.666356

**Authors:** Andreas Zingg, Reto Ritschard, Helen Thut, Gregor Hutter, Heinz Läubli

## Abstract

Chimeric antigen receptor (CAR) T-cell therapy has demonstrated remarkable efficacy in hematological malignancies, but its success in solid tumors remains limited. Conventional CAR-T designs do not integrate into the T-cell receptor (TCR) and rely on non-natural CD3ζ signaling. They thus often suffer from tonic signaling, rapid T-cell exhaustion, and antigen escape due to reduced sensitivity. To address these limitations, we explored alternative CAR architectures that take into account the evolutionary optimized TCR signaling machinery. Specifically, we employed T-cell receptor fusion constructs (TRuCs), where a single-chain variable fragment (scFv) targeting the Sialyl-Thomson-Nouveau antigen (STn) is attached to the CD3ε subunit. We then systematically screened a library of costimulatory domains and their combinations, fused to the TRuC, using a novel in vitro and in vivo screening approach. This screen identified a potent TRuC variant incorporating both CD28 and CD27 costimulatory domains. This optimized construct exhibited increased in vitro and in vivo proliferation and enhanced 4-1BB and IFNγ expression upon repeated antigen stimulation. It further showed improved antitumor efficacy compared to conventional second-generation CAR-T cells in a mouse melanoma model. Moreover, we validated the adaptability of this approach by targeting B7H3 in a sarcoma model. The TRuC with costimulatory domains again outperformed other CAR-variants. Our findings highlight the potential of incorporating combined CD28 and CD27 costimulatory domains into TCR-based CAR architectures to overcome limitations associated with conventional CAR-T therapy and improve efficacy against solid tumors.

## Introduction

Adoptive cell therapy utilizing autologous chimeric antigen receptor (CAR)-T cells has emerged as a promising treatment option for patients with refractory or relapsed leukemia {Guedan, 2019 #399;Du, 2025 #397;Uslu, 2025 #398}. These therapies, which employ second-generation CAR-T cells equipped with a single costimulatory domain, target cancer-associated antigens like CD19 or BCMA {Guedan, 2019 #399}. Despite the improved outcome in hematological malignancies, solid tumors seem more resistant towards CAR-T cell therapy. This is in part driven by the hostile immunosuppressive tumor microenvironment which restricts the access and inhibits the effector function or CAR-T cells. Still, the success of adoptive transfer of tumor-infiltrating lymphocytes (TIL) proves that T-cells are in principle able to completely eradicate advanced, metastatic tumors^1^. TILs show some advantages compared to CAR-T cells: they are polyclonal and specific to genuine cancer antigens (neoantigens) and are more likely to find their way back to the tumor they came from than peripheral blood mononuclear cell (PBMC)-derived T cells used for manufacturing CAR-T cells^2^. In addition, the artificial CAR construct leads to non-native T cell signaling as well as excessive activation upon repeated antigen encounter and is prone to tonic signaling due to aggregation of single-chain variable fragments (scFvs). Elements of 2^nd^ generation CAR-T cells have been optimized to address the most relevant shortcomings with improved performance in preclinical experiments. Alternative costimulatory domains and their combinations were evaluated. The signaling strength was fine-tuned by deleting activating motifs of CD3ζ. Resistance to immunosuppression was improved by knocking-out checkpoint molecules or by expressing decoy-receptors. CARs were programmed to drive overexpression of beneficial cytokines for better survival and proliferation or for recruiting other immune cells. Still, the results of most clinical trials using 2^nd^ or 3^rd^ generation CAR-T cells targeting solid tumor remain below expectations with few exceptions^3, 4^. Therefore, other strategies are needed to leverage the therapeutic potential of CAR-T cells to match the impact of TILs ^5^.

Changes to the overall architecture of CAR-T in which the targeting domain is integrated into the T-cell receptor (TCR) represents an important effort towards this goal as this design more closely resembles the native TCR-signaling. Compared to classic CAR-T cells, tonic signaling was reduced along with improved antigen sensitivity and performance in preclinical *in vivo* experiments. scFvs have been attached to all proteins of the TCR complex^6^ and antibody-derived variable fragments were fused to the constant domains of α/β^7, 8^ or γ/δ^9^ chains. Recent approaches also included the addition of costimulatory domains to α/β chains with promising preclinical and clinical results^10, 11^.

Screening for alternative costimulatory domains has been reported for 2^nd^ and 3^rd^ generation CAR-T cells but only few have been tested in TCR-mimicking configurations.

Here, we describe a novel *in vitro* and *in vivo* screening system for next-generation CAR-T cells. We built and screened a library of selected costimulatory domains and their combinations fused to the CD3ε-based *T-cell receptor fusion construct* (TRuC). The simple design of the TRuC has several advantages over α/β-chain modified constructs. The TRuC integrates into the TCR without disturbing correct pairing of α- and β-chains, a problem that must otherwise be addressed. Engineering and construction of variants is facilitated by the monocistronic CAR expression cassette. The approach of combined *in vivo* and *in vitro* screening allows for systematic investigation and selection of functionally superior CAR-T variants from a large library, not limited to our TRuC-design.

## Results

### Costimulatory domains fused to the TRuC induce improved activation of T cells

We previously reported the generation of Sialyl-Thomson-Nouveau (STn) directed CAR-T cells. Along 2^nd^ generation, CD28- and 4-1BB-based CAR-T cells, we also evaluated an anti-STn-TRuC^12^. Despite the convincing, published data from preclinical experiments^6, 13^, we observed inferior performance of the TruC *in vivo* compared to the CD28-CD3ζ variant (**Supplementary Figure 1**). We hypothesized that the lack of co-stimulation, even when harnessing TCR-like signaling, leads to rapid exhaustion and dysfunctionality. As a proof-of-concept, we fused the two intracellular domains of CD28 and OX40 to the C-terminus of the TRuC and evaluated the effect in a mouse model. CD28-OX40 was chosen due to its reported improved performance as 3^rd^ generation CAR-T cells^14, 15^. When we co-cultured the constructs with STn-expressing HT1080 sarcoma cells for 3 days, we observed that the added costimulatory domains indeed improved the killing capacity (**Supplementary Figure 1b**). At effector-to-target (E:T) ratios below 1:2, both 2^nd^ generation CAR-T cells and TRuC-CD28-OX40 were able to control the growth of tumor cells, while the TRuC failed to do so. We then tested these constructs in a preclinical therapy study with NSG-mice bearing subcutaneous STn-expressing A375 melanoma tumors. When the tumors reached ∼45 mm^3^ (∼14 days), animals were treated with 5 mio CAR-T cells intravenously. Initially, TRuC-CD28-OX40 performed equally well as the classic CD28-CD3ζ CAR-T cells. However, tumor growth-curves started to separate 10 days after treatment and the classic CD28-CD3ζ variant outperformed the co-stimulated TRuC (**Supplementary Figure 1C**). Still, this new variant performed better than the original TRuC and 4-1BB-CD3ζ CAR-T cells.

We concluded from these results that CD28 and OX40 might not be the ideal costimulatory domains and others might be better suited.

### Library construction and sequencing validation of TRuC-variants with costimulatory domains

We wanted to systematically investigate the effect of different costimulatory domains in combination with CD3ε-based CAR-T cells. We constructed additional 10 variants that include commonly used domains such as CD28, 4-1BB and OX40, and combinations thereof. Further, we included other promising, published domains and kept the specificity for STn (**Figure 1a**). Since evaluating 12 different variants in parallel is challenging, especially when multiple PBMC donors should be tested, we decided to pool them and screen the library for outperforming variants. To circumvent the problem of transducing multiple variants into a single T cells, we separately transduced primary human T-cells, tested their surface expression and then pooled the variants (**Figure 1b**). The surface expression was variable but most constructs expressed well, indicating that CD3ε-fusion constructs successfully integrated into the TCR^16^. Only the 4-1BB-containing variants showed a suboptimal expression and separation from non-transduced cells (**Supplementary Figure 2a**).

**Figure 1.**
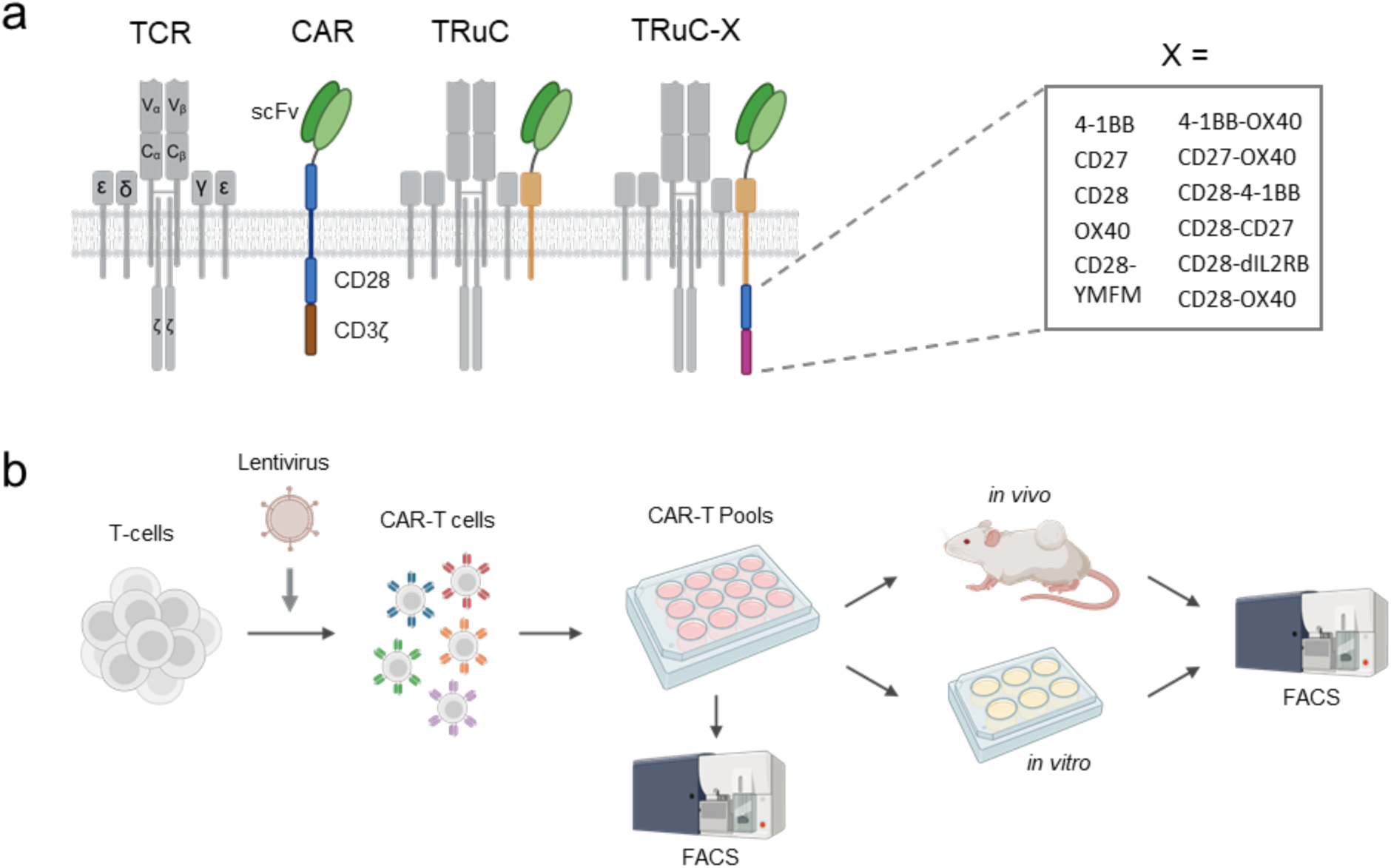
TRuC library construction. **a**, Schematic comparison of TCR, classic second-generation CAR, TRuC and TRuC with various single and dual costimulatory domains. **b,** Overview of the screening process. T-cells are transduced separately with lentivirus, pooled and subjected to *in vitro* and *in vivo* challenge. T-cells with preferred performance were sorted by FACS.

In order to measure the frequency of each variant in the library, we next established an analysis pipeline using next-generation-sequencing. We sorted CD4 and CD8 T-cells separately, whereby CAR-expression was not considered as non-transduced T-cells will not influence the composition of the library. Genomic DNA was isolated and the relevant part of the CAR expression-cassette was amplified by PCR (**Supplementary Figure 2c**). In a secondary PCR, a barcode was introduced to allow pooling of up to 40 samples in a single sequencing-library. As the variants are different in size, we could not exclude an amplification and sequencing bias. To circumvent the issue of determining the actual library composition, we decided to calculate the enrichment factor for each variant by comparing the libraries before and after selection. To further reduce amplification and sequencing variability, all samples were processed in triplicates. Additionally, we analyzed the influence of variable cell numbers used for genomic DNA extraction for 3 libraries. We did not observe considerable variability within triplicates or in regard of cell numbers used as starting material (**Supplementary Figure 2c**). What was observe was a bias towards a higher reported frequency of smaller variants with a single costimulatory domain. Transduction rates and cell counts contradicted this bias though.

### Repetitive stimulation in vitro identifies rapidly proliferating and exhaustion-resistant variants

We next used the generated libraries in repetitive stimulation experiments. We co-cultured the CAR-T cells with three different tumor cell lines that we had previously classified as exhibiting low (HT1080), medium (A375) and high (A549) resistance to CAR-mediated killing. For HT1080 and A375, co-culture was initiated with an equal number of tumor and CAR-T cells. Cell suspensions of tumor cells were added to CAR-T cells after one and two weeks for restimulation (**Figure 2a**). For the A549 cell line, a slightly more challenging setup was chosen. We let the cancer cells adhere for two days and secrete inhibitory molecules into the medium. CAR-T cells were then added and co-cultured for 3-4 days and CAR-T cells were transferred again to conditioned medium with adherent A549 cells. CAR-T cells were challenged three times before sorting or a fourth overnight stimulation. After a week of resting, part of the library was sorted to assess proliferation. The remaining cells were stimulated a fourth time and cells were sorted for 4-1BB and IFNγ expression. We chose these markers as they were previously associated with better outcome when expressed in tumor-infiltrating lymphocytes. 4-1BB and IFNγ should identify non-exhausted T-cells that still show effector function after repeated stimulation. To quantify proliferation, we compared the frequency of variants from baseline with the pool after three stimulations and expansion (**Figure 2b**). The variant frequencies of IFNγ-and 4-1BB-positive cells were compared with their frequencies after repeated stimulation. Overall, we found some variants that showed strong proliferation and retained effector function after repeated stimulation: OX40, CD27, CD28-CD27, CD27-OX40 (**Figure 2b, Supplementary Figure 3a**). Most importantly, these variants were equally effective in CD4 and CD8 T-cells. The data is also consistent with previous *in vivo* data. The TRuC showed poor proliferation and reduced effector function in comparison to the CD28-OX40 variant. The CD28-OX40 variant was further outperformed by the aforementioned costimulatory domains, promising that those will also have better *in vivo* performance.

**Figure 2.**
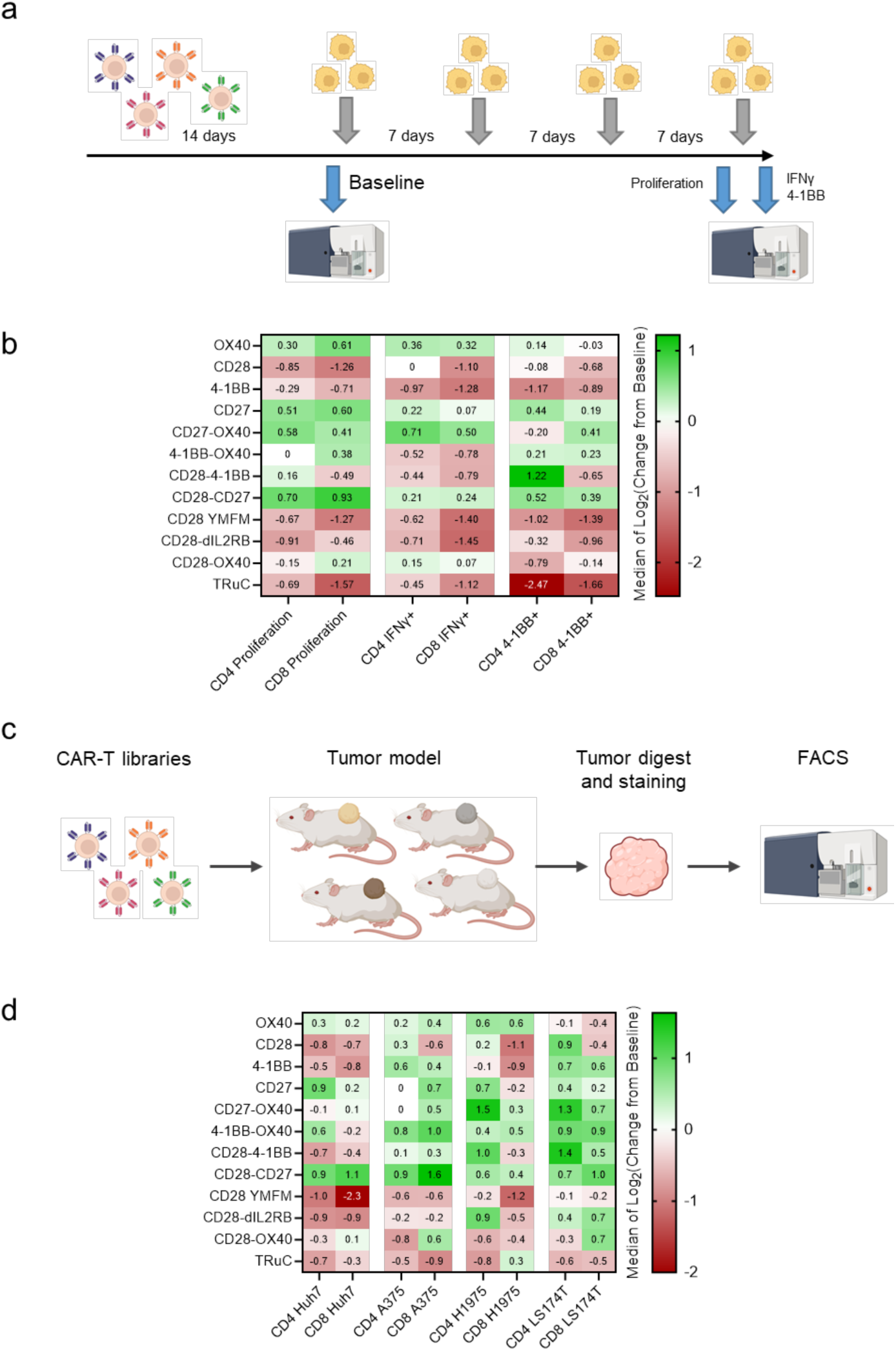
*In vitro* screening of TRuC variants. **a,** Outline of the repetitive stimulation and functional readout. TRuC pools were stimulated weekly by addition of various tumor cells. **b,** Summarizing results of the *in vitro* screen. The median of donors for the same selection conditions is shown. **c,** Outline of the *in vivo* screening. Tumor-bearing mice were injected with TRuC pools and tumors were extracted and digested at endpoint (≤1500 mm^3^). After cryopreservation, CD4 and CD8 T-cells were isolated by FACS. **d,** Summarizing results of the *in vivo* screen. The median of donors for the same tumor model is shown.

### Pooled in vivo screening resulted in similar variants as in the in vitro screening

Weekly stimulation in cell culture medium in presence of IL-2 cannot recapitulate all challenges that CAR-T cells face in a living organism. CAR-T cells need to persist in the absence of stimulation, must migrate to the tumor and should retain prolonged effector function in a hostile tumor microenvironment. The use of immunodeficient mice with a xenograft tumor will also not fully represent the actual situation due to the lack of tumor-infiltrating, immunosuppressing immune cells. Nevertheless, it will provide valuable information on whether the *in vitro* enriched variants will also perform well *in vivo*. We used several tumor models that previously showed infiltration of CAR-T cells. Treated tumors were isolated and digested, and tumor-infiltrating T-cells were sorted by FACS (**Figure 2c**). In total, we analyzed four different tumor cell line in 22 mice, using TRuC CAR-T cells from 9 PBMC donors. Similar variants were enriched as in the *in vitro* repetitive stimulation experiments (**Figure 2d**). Even though both assays enriched for proliferation and resistance towards exhaustion, this result was unexpected as the *in vitro* assay does not select for the ability to migrate to the tumor. This observation indicates that costimulatory molecules do not seem to affect tumor homing and infiltration. The considerable variability between donors emphasizes the need for utilizing multiple donors, in the best case using multiple mice per donor and tumor model. For example, within CD8 CAR-T cells in H1975 tumors, all donors were depleted for the CD28 domain. In contrast, some donors were enriched for CD27 or 4-1BB-OX40 while others were depleted for those variants (**Supplementary Figure 3b**).

### TRuC-CD28-CD27 shows better in vivo tumor control without excessive cytokine secretion

After having identified promising variants, we wanted to confirm our findings and compare them to the classic CD28-CD3ζ CAR. We used again the A375-STn model which we previously used for the proof-of-principle experiment and the *in vivo* screening. When tumors reached 40-50 mm^3^, 4 mio CAR-T cells were injected intravenously. While most variants performed similar to the CD28-CD3ζ CAR and slightly better than the TRuC, the TRuC-CD28-CD27 showed a much better tumor control resulting in prolonged survival (**Figure 3a**). We measured cytokine secretion *in vitro* to check if this characteristic correlated with the outcome of the mouse experiment. However, the results did not reveal marked differences between the variants (**Figure 4b**). Consistent with what we have previously reported, the TRuC only secreted minimal amounts of cytokines compared to the CD28-CD3ζ. Addition of costimulatory domains did not significantly increase IL-2 and TNFα. Only IFNγ secretion was minimally but non-significantly upregulated and the amount was still much less than in the classic CD28-CD3ζ CAR. The observed *in vivo* performance could thus not be explained by cytokine production rate. CAR-T cells secreting less cytokines might even be better tolerated in patients. In addition, cytokines can have both positive and negative effects in tumor control, e.g. IFNγ from immune cells induces PD-L1 expression on tumor cells^17^.

**Figure 3.**
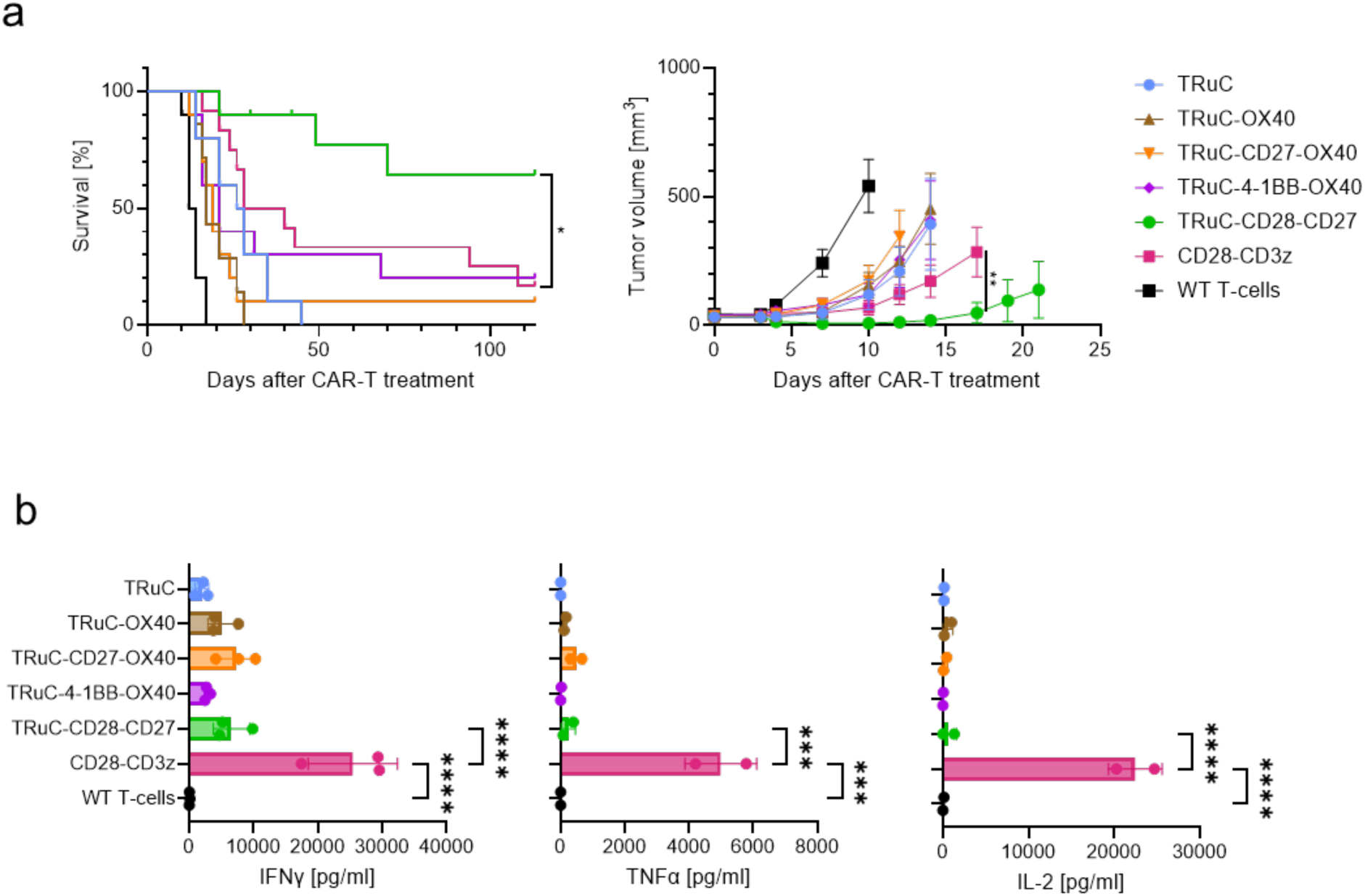
Confirmation of identified constructs in a mouse model. **a**, A375-STn mouse model with selected TRuC-variants. Mice were inoculated with 3.5 mio STn-expressing A375 tumor cells and treated 14 days later with 5 mio CAR-T cells. Statistics show two-way ANOVA performed for day 17 of CD28-CD3ζ and TRuC-CD28-CD27 groups only. WT: untransduced T-cells (wildtype). Survival was compared only for TRuC-CD28-CD27 and CD28-CD3ζ T-cells. Experiment was performed twice using different PBMC donors. 7-12 mice were included in each group. **b**, Analysis of secreted cytokines after 3 days of co-culture with target cells. One-way ANOVA was used for statistical analysis. IFNγ: results from three different PBMC donors, IL-2 and TNFα: results from two different PBMC donors. Comparisons between CD28-CD3ζ and TRuC variants or WT were highly significant (*** or ****), while comparisons within TRuC variants and WT T-cells was not significant.

**Figure 4.**
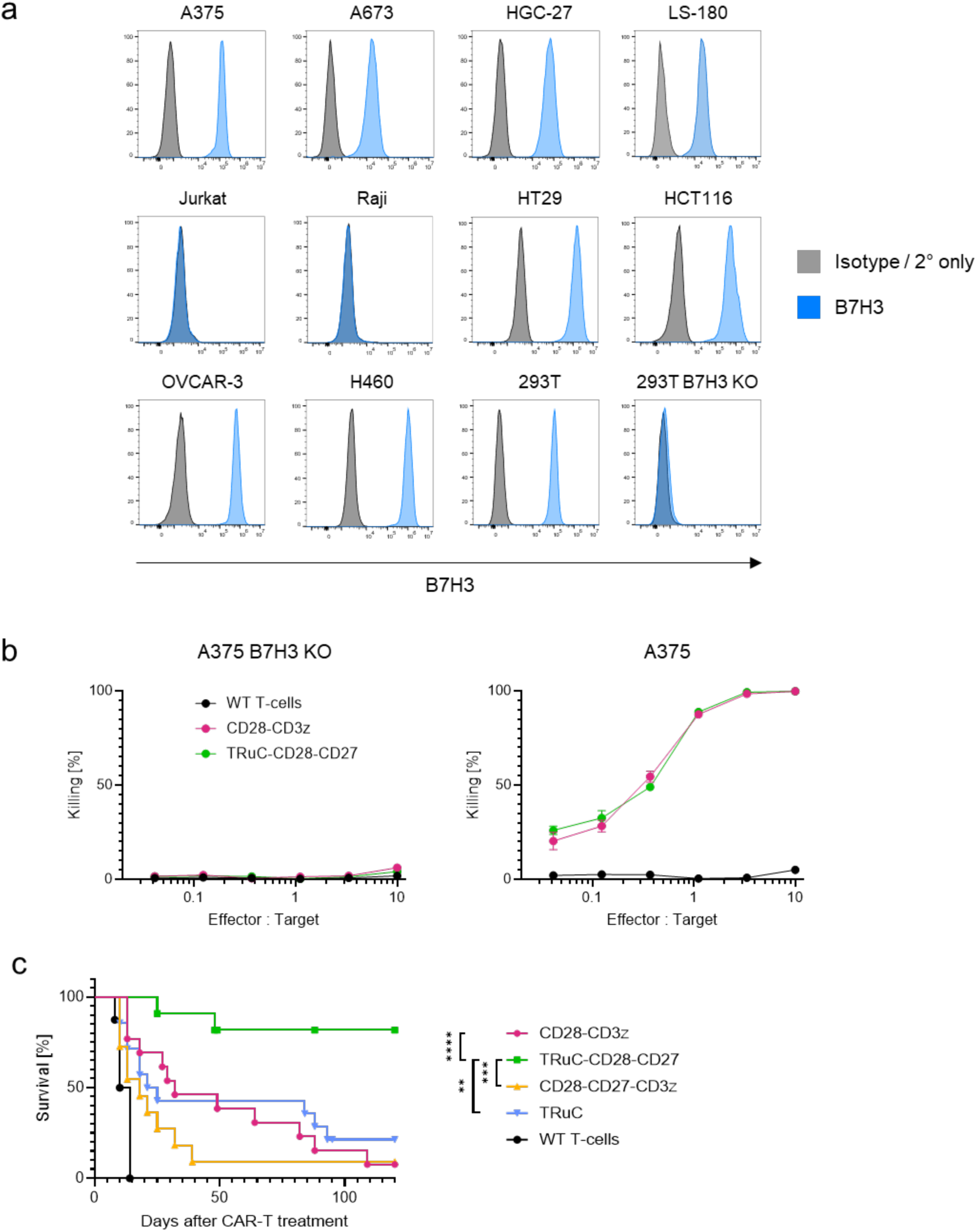
Validation of TRuC-CD28-CD27 variant targeting B7H3. **a**, B7H3 staining of various cancer cell lines with MGA271 (murine version of Enoblituzumab). **b**, Cytotoxicity assays show equal *in vitro* potency of CD28-CD3ζ and TRuC-CD28-CD27 targeting B7H3 after 5 days of co-culture. B7H3-targeted CARs do not show off-target toxicity for the A375-B7H3 knock-out cell line. **c**, Survival curve of mice implanted with 5 mio A673 sarcoma cells and treated 6-7 days later with 5 mio CAR-T cells targeting B7H3. Tumor-free mice were re-challenged 60-70 days after the first CAR-T treatment with 5 mio A673 tumor cells.

### B7H3-targeting TRuC-CD28-CD27 retains its improved tumor control capabilities

The obtained results suggested that the TRuC-CD28-CD27 is the most promising design. To generally validate the platform and to evaluate whether this design is also efficient with other targeting domains, we explored B7H3 as promising clinical target. The expression of B7H3 is high in the majority of tumors^18, 19^ but limited in healthy tissue. Numerous clinical trials using CAR-T cells^20, 21^ and other treatment strategies targeting B7H3^22, 23^ are ongoing.

We constructed an scFv derived from Enoblituzumab^24^ containing the Withlow linker^25^ and fused it to the CD28-CD3ζ CAR domains. We first evaluated the specificity of this new combination by co-culturing CAR-T cells with various cancer cell lines. Cell lines were all highly positive for B7H3, except for the HEK293T knock-out cell line (**Figure 4a**). Intracellular cytokine staining and 4-1BB upregulation were assessed by flow-cytometry. As expected, B7H3-targeted CD28-CD3ζ CAR-T cells were not activated by the knock-out cell line but expressed IL-2, IFNγ, TNFα and 4-1BB upon target recognition (**Supplementary Figure 4a**).

We further generated B7H3-targeting TRuC variants either without or with CD28-CD27 costimulatory domains. Additionally, we included a third-generation CD28-CD27-CD3ζ variant to evaluate if the addition of the CD27 costimulatory domain improved the function in the context of a conventional CAR constructs as well.

For the CD28-CD3ζ and TRuC-CD28-CD27 variants, we performed killing assays with the unmodified A375 cell line as well as with the B7H3 knock-out cell line. No differences in cytotoxicity between the variants after 5 days of co-incubation (**Figure 4b**) was observed. B7H3-targeting CARs efficiently killed the B7H3-expressing A375 cancer cells even at low effector-to-target ratios but spared the knock-out cell line which indicates little tonic signaling and is consistent with low cytokine expression in absence of target cells. We further evaluated the serial killing ability in a repeated stimulation assay. CD28-CD3ζ, TRuC and TRuC-CD28-CD27 CAR-T cells were transferred twice a week onto a new plate with pre-seeded A673 cancer cells. 3 days after transfer, killing efficiency was measured and CAR-T cells were transferred to a new plate with tumor cells. A total of four rounds of stimulation were performed. Surprisingly, CD28-CD3ζ CAR-T cells performed better than the TRuC variants (**Supplementary Figure 4b**).

We still proceeded to a subcutaneous Ewing sarcoma mouse model using the A673 cell line. NSG mice were subcutaneously injected with 5 mio A673 cells and treated with 5 mio CAR-T cells 6-7 days later once a measurable tumor has formed (∼35 mm^3^). Mice that were tumor-free 60-70 days after initial treatment were re-challenged with 5 mio A673 cells. Mice that were treated with TRuC-CD28-CD27 T-cells were completely resistant towards re-challenge and rapidly cleared the engrafted tumor cells while half of the TRuC T-cell treated mice relapsed. The TRuC-CD28-CD27 T-cell clearly outperformed all other variants regarding tumor control, persistence and survival (**Figure 5c**, individual tumor growth curves: **Supplementary Figure 4c**). The contradicting results from repeated stimulation and *in vivo* results again highlight the necessity of mouse models and the short-comings of pure *in vitro* analysis.

These results demonstrate that our *in vitro* and *in vivo* screening strategy was able to select an improved TRuC construct outperforming classical CAR constructs in preclinical models.

## Discussion

The development of CAR-T therapies represents an important milestone in the treatment of hematological malignancies. The success of TIL therapy in metastatic melanoma demonstrates that adoptive cell therapy is in principle feasible and potent also for solid tumors. However, CAR-T cell therapy so far showed limited success for the treatment of solid malignancies. Successful T-cell therapy in solid tumors depends on numerous factors including efficient tumor infiltration and persistence in a hostile environment. A major limitation of traditional 2^nd^ generation CAR constructs is the unregulated, non-native signaling resulting from the artificial architecture. This inadequate signaling can accelerate exhaustion and limit the effector function of CAR-T cells^26^.

An emerging strategy is the fusion of the targeting protein to domains of the TCR. The TRuC approach is particularly elegant as the CAR construct is linked to the CD3ε domain which efficiently integrates into the TCR without the need for engineering the TRAC/TRBC locus. Despite previously published, promising results, our STn-targeted TRuC T-cells exhibited inferior *in vivo* therapeutic activity compared to the classic CD28-CD3ζ CAR-T cells in preclinical models of cancer. The addition of CD28-OX40 costimulatory domains to the TRuC improved its performance but was still not better than the CD28-CD3ζ CAR variant. This finding encouraged our hypothesis but also highlighted the need to optimize the costimulatory domains of the TRuC. To enable systematic exploration of a significant number of costimulatory domains in a pooled library comprising 12 variants, we developed a combined *in vitro* and *in vivo* screening platform. The *in vitro* selection is based on several readouts which were previously shown to be associated with increased *in vivo* performance such as proliferation, 4-1BB upregulation and IFNγ secretion after repeated stimulation. During the *in vivo* selection, the enrichment of TRuC variants in the tumors following administration of a sub-therapeutic dose of the CAR-T library was evaluated after tumor outgrowth. An important strength of this approach is that a relatively large number of costimulatory domains and combinations thereof could be evaluated in parallel in different mouse models and with PBMCs derived from different donors. The latter was found to be especially important as considerable heterogeneity was observed for certain models. However, the necessity of the *in vivo* screen could not be confirmed in our experimental setup as it enriched the same variants as the *in vitro* screening. It cannot be excluded though that a larger library or an alternative focus such as screening for different binding domains could benefit from *in vivo* screening models. The approach described in this manuscript may be generally applicable to other CAR architectures and not only for the TRuC. Our library of 12 variants was small compared to other screening approaches but showed the feasibility of an *in vivo* screen. More dedicated cloning strategies allow for building larger libraries and increase the chance of finding promising variants. The inclusion of genetic barcodes would additionally facilitate the library preparation and analysis by next-generation sequencing^27^.

Both the *in vitro* and *in vivo* selections consistently identified the TRuC-CD28-CD27 as most promising architecture. This finding supports the hypothesis that the costimulatory domain is mostly important for persistence and effector function in the tumor but may be secondary for tumor homing as all variants were readily found in the tumor. In addition, the differences observed between the various constructs highlight the importance of optimizing the combination of costimulatory domains in the development of CAR-T cell therapies as neither CD28 nor CD27 alone were either not enriched or not effective. The TRuC-CD28-CD27 variant showed less cytokine secretion *in vitro* compared to the classic CD28-CD3ζ CAR-T cells but exhibited more potent anti-tumor activity *in vivo.* This might indicate that the artificial signaling in CD28-CD3ζ CAR-T cells drives overstimulation which might be detrimental *in vivo* while *in vitro* rather the opposite was observed. It also emphasizes the importance of *in vivo* evaluation of CAR-T constructs. A possible limitation is the use of non-MHC knock-out mice. We cannot fully exclude the influence of mispairing of human TCRs and mouse MHCs. We observed severe GvHD, assumed by weight loss, only in few of the TRuC-CD28-CD27 treated mice that had cleared the tumors. The clearance of the A375-STn tumor, however, was mediated by antigen-specific CAR-activation as the tumor cells did not express MHC-I molecules (knock-out of beta-2-microglobulin).

To validate the general applicability of this approach and the improved performance of the TRuC-CD28-CD27 variant, we evaluated B7H3 as another target of high clinical importance. The CD28-CD3ζ and TRuC-CD28-CD27 variants showed high selectivity towards B7H3-positive tumor cells in an *in vitro* cytotoxicity assay but did not differ otherwise. For the *in vivo* validation, we additionally included a classic third-generation CAR containing CD28 and CD27 as costimulatory domains to distinguish the contribution of those to both the TRuC and CD3ζ architectures. Mice bearing subcutaneous sarcoma tumors that were treated with the TRuC-CD28-CD27 T-cells rapidly cleared the tumors and were entirely resistant towards repeated re-challenge. 9 out of 11 mice remained tumor-free until the end of the experiment (120 days) without showing side-effects such as weight-loss.

In conclusion, we described a novel *in vitro* and *in vivo* screening platform which enables an efficient systematic investigation of costimulatory domains for CAR constructs. We discovered a new combination of costimulatory domains for TRuC-type CAR constructs and demonstrate that this approach is portable to other targeting domains.

## Materials and Methods

### Plasmid construction

All plasmids were planned *in silico* with Geneious Prime (V2024.0.3, RRID:SCR_010519). DNA sequences were acquired from publications, patents^28^, Uniprot (RRID:SCR_002380) or the NCBI database. Protein-coding regions were codon-optimized with Genscript condon optimizer (RRID:SCR_002891), ordered as DNA fragments from Genscript or Twist Bioscience and cloned into lentiviral plasmid backbones (pLV, VectorBuilder) under the control of the EFS (EF1α-short) promoter.

### Flow cytometry

Cells were analyzed on CytoFLEX flow cytometers (Beckman Coulter, RRID:SCR_025067). All stainings were performed in PBS with 1% FBS and 2 mM EDTA with incubation steps at 4°C for 30 min. DAPI was used as a live-dead marker if not otherwise mentioned. CAR-T cells were stained with recombinantly expressed anti-Whitlow-linker antibody “Clone 16” (Biointron; Patent: WO2018013563A1 ^29^) and goat-anti-rabbit-PE antibody (Invitrogen, Cat: P-2771MP). Cell lines were analyzed for B7H3 expression by recombinantly expressed murine MGA271^30^. Data was analyzed with FlowJo (V10.10.0, RRID:SCR_008520). All staining panels are listed in **Supplementary Data 5**.

### Cell Culture

A375-Luc2 were acquired from the American Type Culture Collection (ATCC, RRID:CVCL_UR32, received in April 2022) in which the β2m gene was knocked-out (kind gift from S. Tundo, Cancer Immunology Lab, University of Basel). The HT1080 (RRID:CVCL_0317) and Huh7 cell lines (RRID:CVCL_0336) were a kind gift from F. Stenner (University Hospital Basel, received in October 2022) and authenticated by STR analysis (March 2025, Microsynth). H1975 (RRID:CVCL_WV99), LS174T (RRID:CVCL_1384) and A673 (RRID:CVCL_0080) were acquired from Cytion. A549 cell line (RRID:CVCL_0023) was acquired from the DSMZ. Lenti-X HEK293T cells were acquired from TakaraBio (RRID:CVCL_4401, received in November 2022). A375, HT1080, H1975, Huh7 and LS174T were grown in RPMI medium. A673 and HEK293T were grown in DMEM medium. RPMI and DMEM media were supplemented with 10% FBS, 1% penicillin/streptomycin, 1% non-essential amino acids and 0.25 ug/ml amphotericin B. Cell lines were regularly tested for mycoplasma contamination. Upon receipt, cell banks were made (1-2 passages) and stored in liquid nitrogen. Cell lines were kept in culture for a maximum of 4 week, afterwards a new vial from the working or master cell bank was thawed. A375, H1975, Huh7 and LS174T were transduced with a ST6GALNAC1 encoding lentivirus as previously described^31^ to enforce expression of STn.

### Knock-out of B7H3 in A375 and Hek293T cells

1.5 ul sgRNA (Genscript, 100 uM, GTGCAGCCCTGGGAGCACTG) and 0.81 ul Cas9 protein (Genscript, 62 uM) were incubated for 20 min at room temperature at the molar ratio of 3:1 to form ribonucleoprotein particles (RNP). 5 mio cells were washed twice with electroporation buffer (Maxcyte) and resuspended in 45 ul electroporation buffer. Cell suspension and RNPs were mixed right before electroporation (program: “Optimisation 4”). Cells were recovered for 4 hours in RPMI or DMEM+10%FBS without antibiotics. Knock-out efficiency was assessed by flow cytometry 7 days after electroporation. Cells were sorted twice for absence of B7H3 expression.

### Lentivirus production

16 hours before transfection, 15 mio Lenti-X 293T cells were seeded in 18 ml complete DMEM medium into 15-cm culture dishes. For the transfection-mix, 2.5 ug pMD2.G (RRID:Addgene_12259), 4.7 ug pCMVR8.74 (RRID:Addgene_22036) and 7.2 ug pLV transfer vector were mixed in 1.8 ml jetOPTIMUS buffer (Polyplus, Cat: 101000025). 18 ul jetOPTIMUS was added and incubated for 10 min before adding the transfection-mix to the cells. Medium was exchanged after 5-6 hours and lentiviral particles were collected after 24 and 48 hours after medium exchange. The pooled supernatant was concentrated with 4X in-house made PEG-8000 solution and resuspended in 1 ml Prime XV serum-free T-cell medium (Irvine Scientific, Cat: 91154). Aliquots were stored at -80°C.

### Generation of CAR-T cells

Peripheral blood mononuclear cells (PBMCs) were isolated from buffy coats (Blutspendezentrum SRK beider Basel) by density centrifugation with Lymphoprep using SepMate tubes (Stemcell Technologies, Cat: 18061, Cat: 85460). Total PBMCs were then activated (10 mio PBMC/ml) with CD2/CD3/CD28 Immunocult T-cell activator (Stemcell Technologies, Cat: 10990) in Prime XV chemically-defined T-cell medium, supplemented with IL-15 (10 ng/ml, Peprotech, 200-15-50UG) and 1% penicillin/streptomycin (Thermofisher). 2 days after activation, cells were isolated and resuspended in 2.5 ml medium containing 0.5 mg/ml synperonic F108 (Sigma, Cat: 07579-250G-F) and 0.125 uM BX795 (MCE, Cat: HY-10514). 100 ul of concentrated lentivirus was added and cells were cultured in 6-well GRex plates (WilsonWolf, Cat: 80240M). After 48 hours, wells were filled up with Prime XV supplemented with 10 ng/ml IL-15 and 1% penicillin/streptomycin and expanded for 14 days. For untransduced T-cells (WT T-cells), instead of lentivirus, Immunocult was added again but were treated otherwise the same. Medium was exchanged every 5-7 days. *In vitro* experiments were performed within 2-3 weeks after production start, animals were treated within 2 weeks after production start.

For transduction of the library variants, transduction was scaled down to 1 ml. Single variants were pooled after 7 days and expanded in 6-well GRex plates.

### Cytotoxicity assay

10’000 GFP+ target or non-target cells were plated in 96-well flat bottom plates in complete RPMI medium without cytokines. 24 hours later, CAR-T cells were 3-fold serially diluted and added to target cells, resulting in an effective 10:1 CAR-to-target-cell ratio for the highest dilution. An equivalent number of untransduced T-cells was added as control. The occupied area-fraction of GFP+ target cells was measured by fluorescence microscopy (Nikon Eclipse Ti2, RRID:SCR_021068) every 24 hours after addition of T-cells for 5 days. For serial killing experiments, 10’000 CAR-T cells in 50 μl were added to 10’000 A673-GFP target cells that were plated 24 hours before in 100 ul RPMI. 72 hours later, killing efficiency was assessed by fluorescence microscopy. CAR-T cells were then resuspended, centrifuged, resuspended in 50 ul fresh RPMI medium and again added to A673-GFP cells that were plated the day before. This procedure was repeated another 3 times.

### Mouse models

NSG mice (NOD.Cg-Prkdcscid Il2rgtm1Wjl/SzJ, RRID:IMSR_JAX:005557) were bred in the animal facility of the Department of Biomedicine (University Hospital Basel) and experiments were performed under approval of the local Ethical Committee (Basel Stadt) and complied with ethical regulations. License number: 3036_34231.

For *in vivo* tumor growth experiments, tumor cells were injected subcutaneously in the right flank in a volume of 100 μl DMEM without phenol red. The following conditions were used for *in vivo* library screening: H1975-STn: 2 mio, treatment after 21 days; A375-STn: 3.5mio, treatment after 15 days; LS174T-STn: 2.5 mio, treatment after 15 days; Huh7-STn: 2 mio, treatment after 15 days. The following conditions were used for CAR-T experiment with single variants: A375-STn: 3.5 mio, treatment after 14 days with 4 mio CAR-T cells; A673: 5 mio, treatment after 6-7 days with 5 mio CAR-T cells. Tumor size was measured three times a week with a caliper (volume = 0.5*width^2^*length) and mice were sacrificed before tumor volume exceeded 1500 mm^3^ or when significant discomfort was observed (e.g. eye infection, weight loss ≥15%).

### Repeated stimulation

For each of three donors, 1 mio TRuC T-cells were mixed with 1 mio HT1080-STn or 1 mio A375-STn cancer cells in 8 ml assay medium (AIM-V, 5% human serum, 200 IU/ml Proleukin) in 24-well GRex plates. One and two weeks later, again 1 mio target cells were added. Medium was changed twice per week. T-cells were left to recover for 2 weeks after the last stimulation.

In a different setting, 200’000 A549-STn cells were cultured for 2 days in 3 ml assay medium in 6-well plates until confluent. 1 mio TRuC T-cells were added and co-cultured for 3-4 days. After this, half of the T-cells were isolated and added to a new co-culture with previously plated A549-STn cancer cells. This process was repeated again. After recovery for two weeks in 24-well GRex plates, TRuC T-cells were subjected to a fourth stimulation.

After co-cultures of HT1080-STn and A375-STn, a fraction of the library was used to evaluate the re-stimulation capacity (IFNγ and 4-1BB expression) after a fourth stimulation. Donor 1 was unresponsive after co-culture with A375-STn cells, while Donor 2 and 3 did not show exhaustion when co-cultured with HT1080-STn target cells. The fourth stimulation was performed using the same cell line as before, using 2 mio T-cells and 1 mio target cells.

Stimulation was performed overnight. For IFNγ intracellular cytokine staining, brefeldin A was added to cultures. TRuC T-cells were sorted the next day using the Cytoflex SRT for CD4 and CD8 cells separately. Maximally 200’000 T-cell were sorted per condition.

Donor 1 was sorted for: proliferation after A549-STn, HT1080-STn and A375-STn co-culture, 4-1BB upregulation after HT1080 co-culture and IFNγ upregulation after HT1080 and A549 co-culture. Donor 2 and 3 were sorted for: proliferation after A549-STn, HT1080-STn and A375-STn co-culture, 4-1BB upregulation after A375-STn co-culture and IFNγ upregulation after A375-STn and A549-STn co-culture.

### Tumor sample digestion and sorting

Tumors were first minced manually and mixed with 5X volume of *in-house* made tumor digestion buffer (45% DMEM, 5% FBS, 50% accutase, collagenase IV (1 mg/ml), hyaluronidase (1 mg/ml) and DNAse I (type IV, 100 units/ml). Digest mix was then incubated for 40 min in shaking incubator (37°C, 50 rpm) and subsequently filtered (100 um). Cells were washed twice with DPBS and frozen in FBS containing 10% DMSO. After cryopreservation, tumor digests were stained with anti-CD45-BV510, anti-CD4-PE and CD8-FITC and sorted for CD4 and CD8 CD45+ cells using the BD Melody cell sorter. DAPI was used as a live/dead marker.

### NGS library preparation

Sorted cells were resuspended in 50 μl KAPA Express Extract (Roche) DNA extraction buffer. Cell suspensions were divided into triplicates and then incubated for 20 min at 75°C before heat-inactivation at 95°C for 5 min. Primary PCR was performed with the following conditions: 5 μl DNA extract, 1.25 μl forward primer (10 μM, CAGAGGGGCCAGAACAAGG), 1.25 μl reverse primer (10 μM, CAAAGGCATTAAAGCAGCGTATCC), 5 μl water, 12.5 μl KAPA2G Fast Genotyping Mix. Cycling program: 3 min @ 95°C, 46 cycles of: 15 s @ 95°C / 15 s @ 60°C / 20 s @ 72°C, 2 min @ 72°C. The resulting PCR product was dilute 100-fold with water for the secondary PCR that was performed with the following conditions: 2 μl diluted primary PCR product, 2.5 μl barcoded forward primer (5 μM, NNNNNNNNNNCAGAAAAGGTCAGCGAGACC), 1.25 μl reverse primer (10 μM, CGTATCCACATAGCGTAAAAGGAGCAAC), 6.75 μl water, 12.5 μl KAPA2G Fast Genotyping Mix. Cycling program: 3 min @ 95°C, 5 cycles of: 15 s @ 95°C / 10 s @ 65°C / 10 s @ 72°C, 33 cycles of: 15 s @ 95°C / 20 s @ 72°C, 2 min @ 72°C. PCR product was purified using 1X SPRIselect magnetic beads according to manual and dissolved in TRIS buffer. DNA was quantified by Nanodrop and single PCR products were pooled accordingly. The pooled libraries were sequenced by Eurofins Genomics, Konstanz using NovaSeq 150 bp paired-end sequencing technology. The resulting reads were processed and analyzed using Geneious Prime 2024.0.7. In brief, reads were separated by barcode and mapped to the sequences of the TRuC variants. The relative frequency of each variant within a library was calculated based on the read-counts and was compared to the relevant control. The summarizing values were calculated as following: median of (log2(relative frequency of variant after stimulation) – log2(relative frequency of variant in baseline control)). Results were plotted using GraphPad Prism 10.3.0.

### Intracellular cytokine analysis

100’000 CAR-T cells were mixed with 50’000 target or non-target cells in RPMI + 10% FBS and 5 ug/ml brefeldin A (Biolegend). Co-cultures were incubated for 5 hours in a humidified incubator. Cells were stained with Zombie violet (Biolegend) as live/dead marker, CD4-FITC and CD8-BV510. After washing, cells were fixed with 100 μl IC Fixation Buffer and subsequently permeabilized (eBioscience). Blocking and staining was performed in permeabilization buffer containing 10% FBS. IFNγ-BB700 and TNFα-APC and IL-2-PE were used at 1:500 and staining was performed overnight. Cytokine expression was normalized to CAR expression.

### Analysis of secreted cytokines

100’000 CAR-T cells were mixed with 50’000 A375-STn target cells in AIM-V medium + 10% FBS. Supernatant was harvested after 3 days of co-culture and cytokine secretion was analyzed using the Lumit human IFNγ Immunoassay (Promega) according to the manual. Analysis was performed in duplicate after diluting the supernatant 4-fold with assay medium.

### Statistics

GraphPad Prism version 10.1.2 (V10.1.2, RRID:SCR_002798) was used for statistical analysis and for visualization of data.

The Log-rank (Mantel-Cox) method was used to compare survival curves of mice treated with CAR-T cells. Tumor volume was compared at the timepoint where the first mouse had to be sacrificed. Tumor volumes were log-transformed after zeros were replaced with 1’s. Two-way ANOVA was performed followed by Tukey’s multiple comparisons test.

Expression frequencies for >2 groups were compared using one-way ANOVA with Tukey’s correction for multiple comparisons. If not otherwise stated, the mean and standard deviation is shown. P-values were assigned as followed: P≤0.05 (*), P≤0.01 (**), P≤0.001: (***), P≤0.0001 (****).

## Authors contributions

A.Z. and H.L. conceived the project and wrote the manuscript. A.Z. planned, performed, and analyzed the experiments. R.R. and H.T. performed experiments. H.L. and G.H. supervised the work and provided reagents.

## Acknowledgments

This work was supported by funding from the Swiss National Science Foundation (SNSF Nr. 310030_184720/1 to H.L.).

## Supplementary Data

**Supplementary Figure 1.**
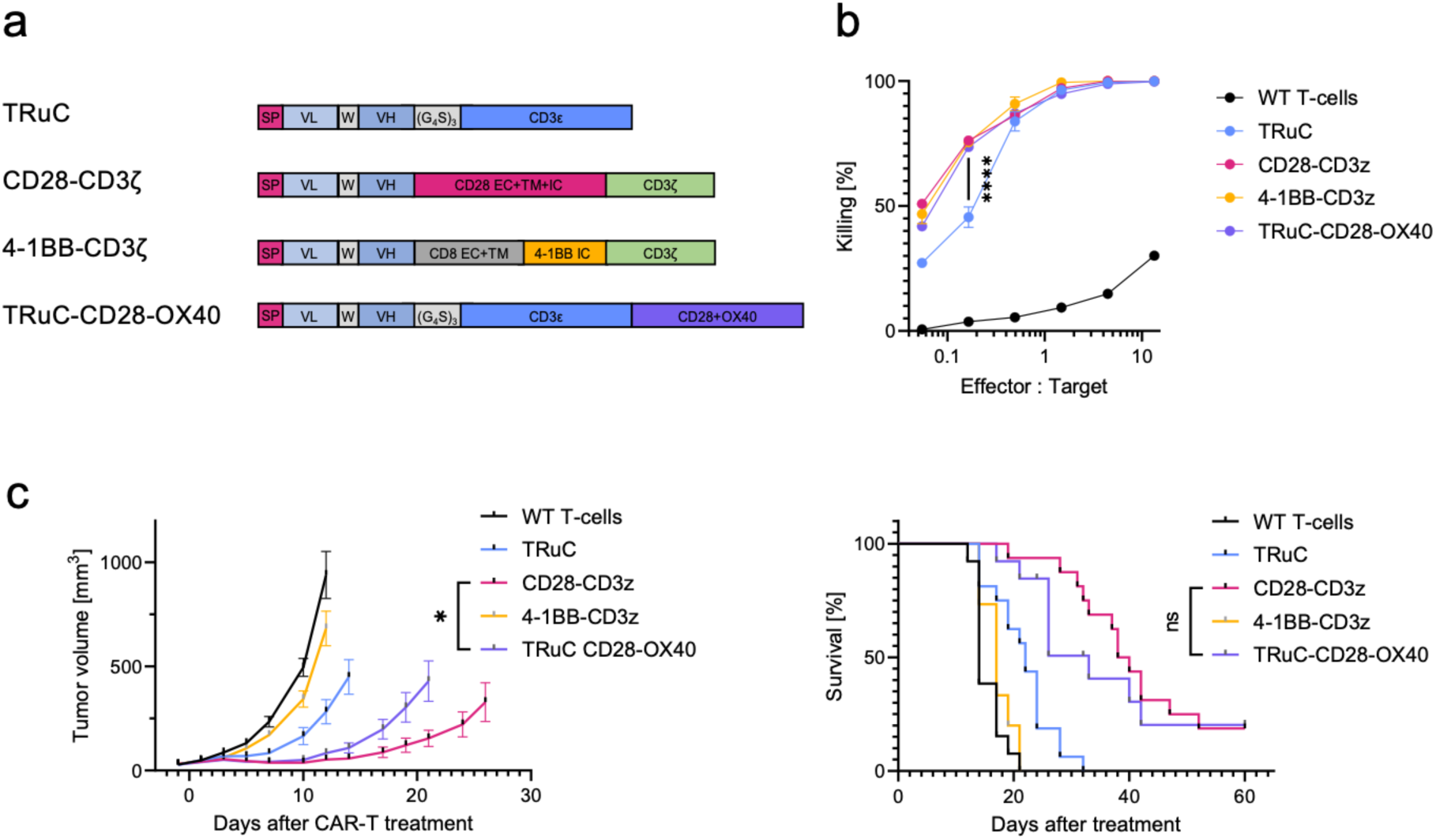
STn-targeted TRuC variants are active *in vitro* and *in vivo*. **a**, SP: signal peptide, VL: variable light chain, W: Whitlow-linker, VH: variable heavy chain, EC: extracellular domain, TM: transmembrane domain, IC: intracellular domain. **b**, Cytotoxicity assay using HT1080-STn target cells. Result after 3 days of co-culture. One-way ANOVA reveals reduced killing of the TRuC, while adding costimulatory domains improves the cytotoxicity. **c**, Mouse model using A375-STn cancer cells. Two-way ANOVA for day 21 of tumor growth, performed only for CD28-CD3ζ and TRuC-CD28-OX40. Survival analysis by log-rank method. Tumor-free mice that were sacrificed during the experiment due to other reasons were blinded.

**Supplementary Figure 2.**
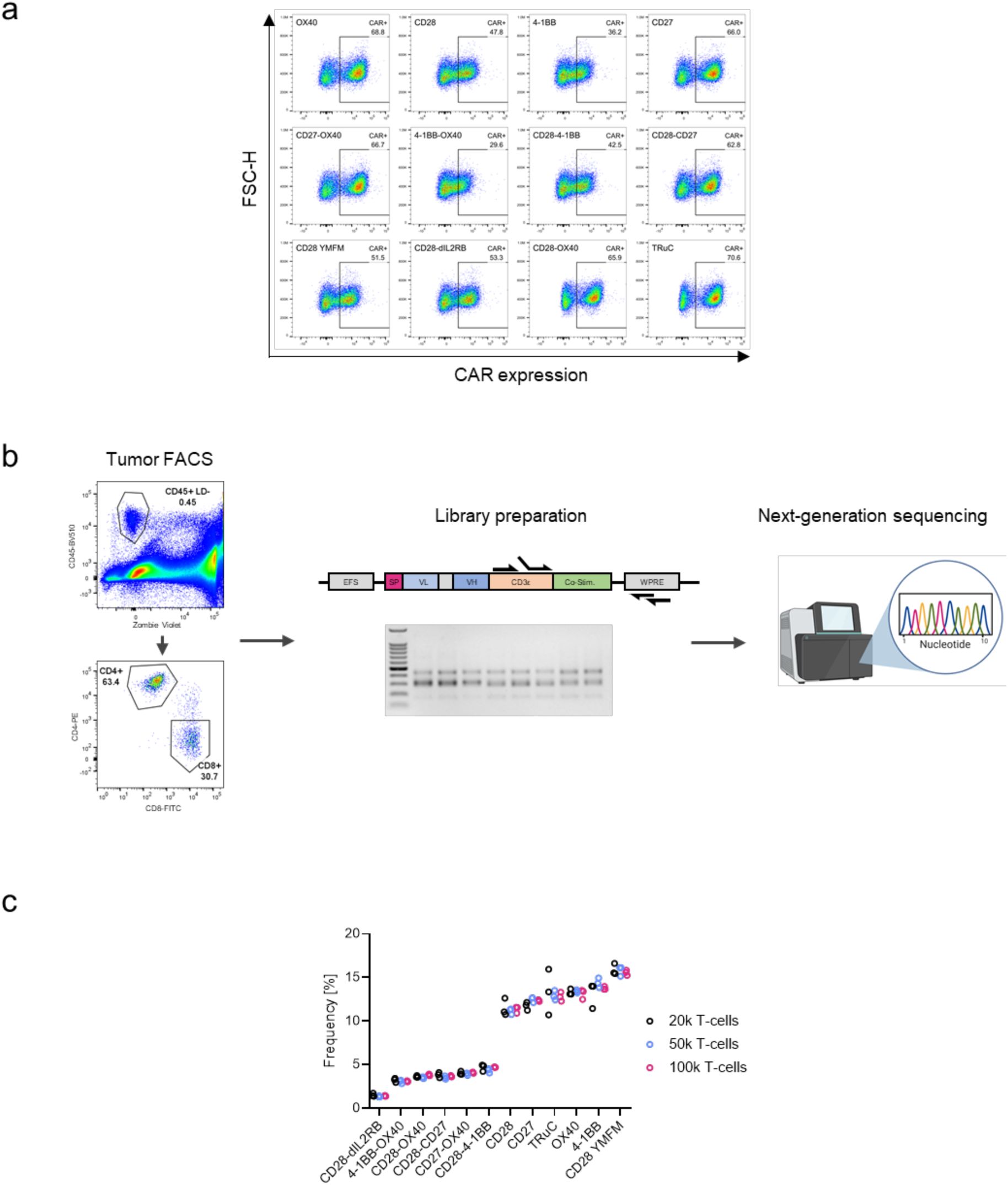
**a** Expression of TRuC variants on primary human T-cells. **b,** Example of tumor digest used for sorting of tumor-infiltrating T-cells. Relevant part of the TRuC DNA was amplified from sorted T-cells in two PCR steps and sequenced **c,** Influence of the number of T-cell used for library preparation and variance of technical triplicates after sequencing.

**Supplementary Figure 3.**
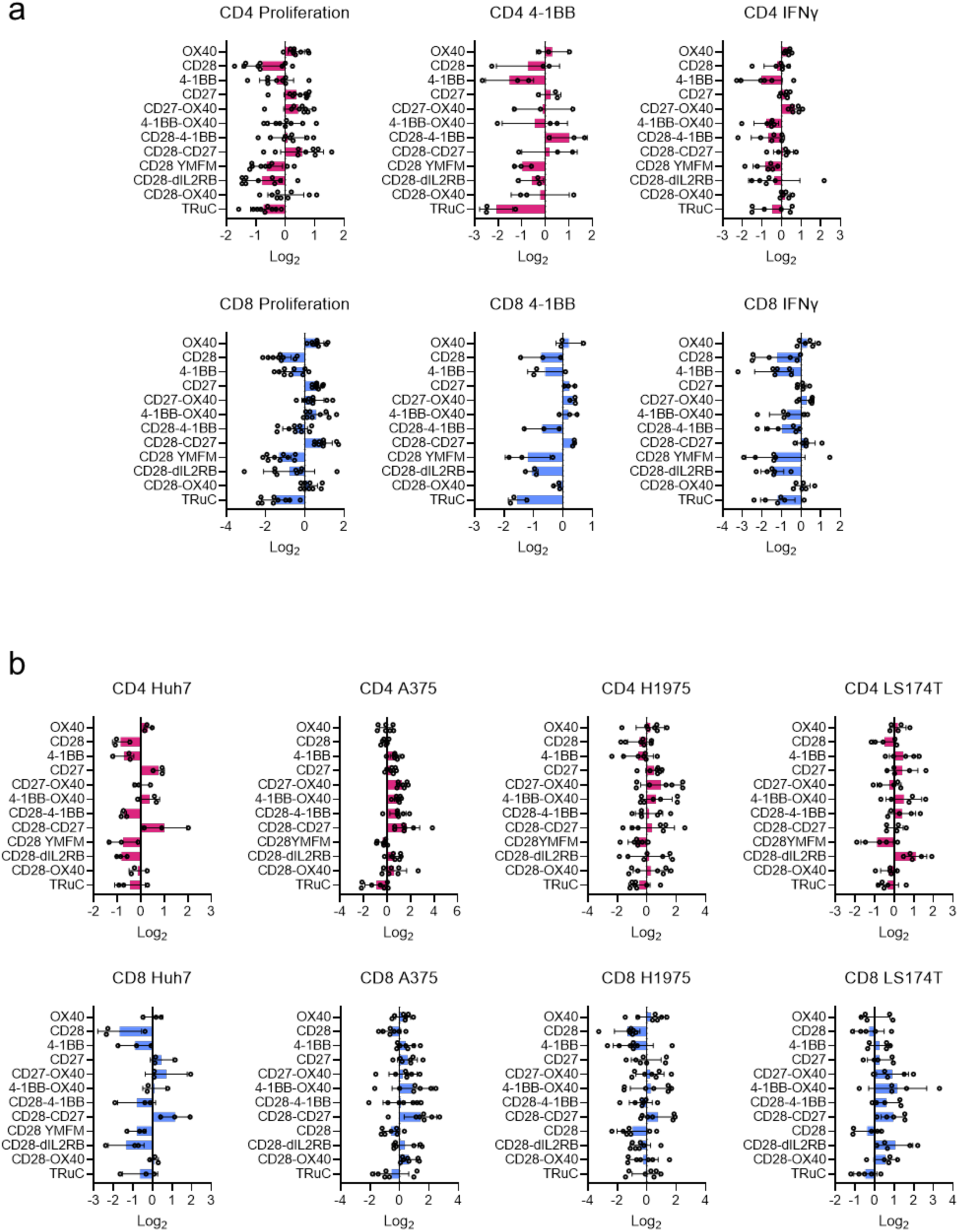
**a**, Detailed results for the *in vitro* screening. Each dot represents one donor under one condition. 3 donors were used for *in vitro* screening. **b**, Detailed results for the *in vivo* screening. Each dot represents a different donor within a row.

**Supplementary Figure 4.**
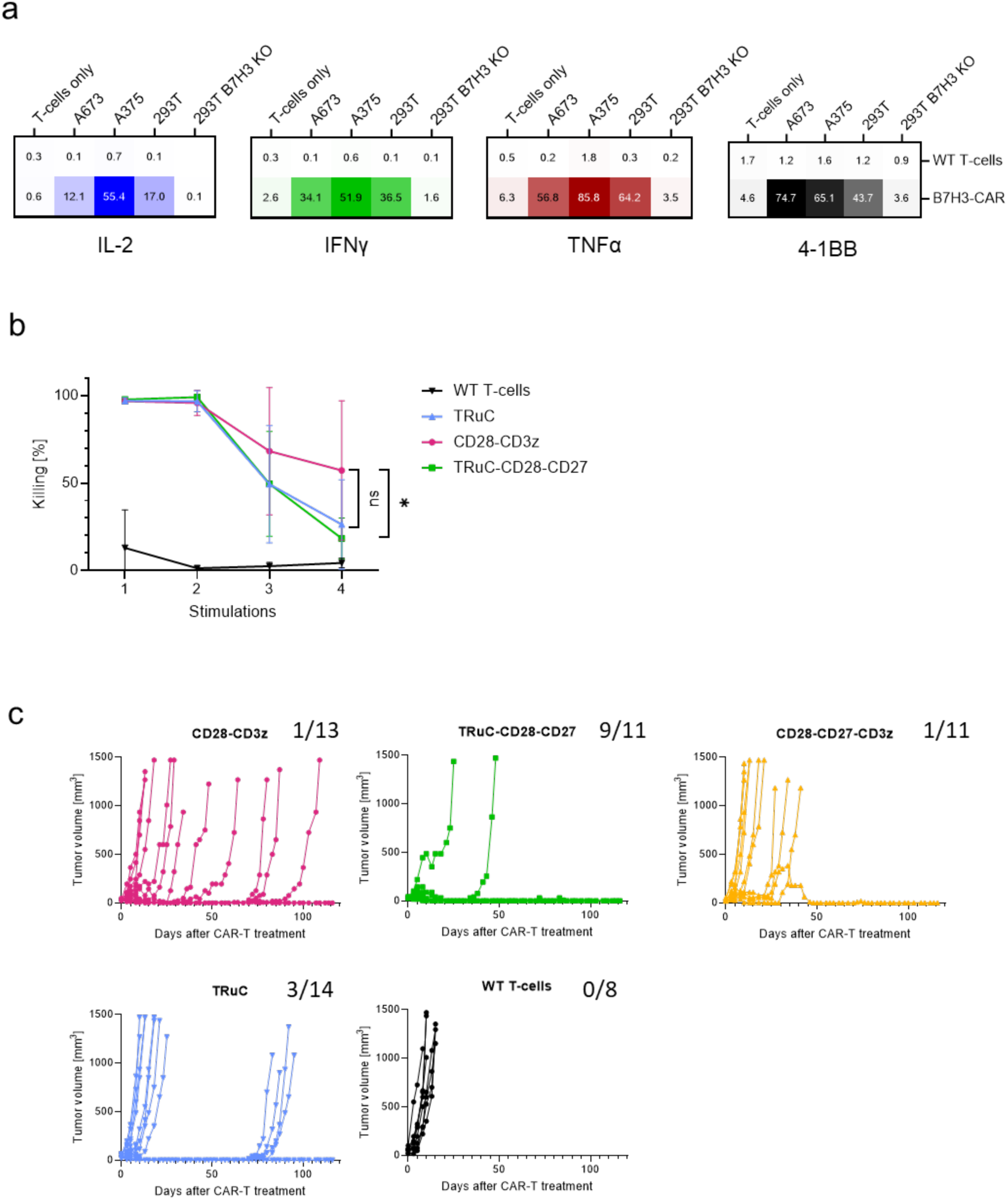
**a,** Intracellular cytokine expression frequency of CAR variants after co-culture with tumor cells, normalized to CAR expression frequency. Results from one PBMC donor. **b**, Cytotoxicity after multiple rounds of stimulation with target cells. c, Individual tumor growth curved of B7H3-targeted CAR-T cells in the A673 sarcoma model. Numbers indicate total number of mice and how many survived until the end of the experiment (120 days).

---

| Intracellular Cytokine Staining |  |  |  |  |  |  |
| --- | --- | --- | --- | --- | --- | --- |
| Marker | Clone | Fluorophore | Dilution | Supplier | Catalog Nr. | RR_ID |
| Live/Dead | Zombie Violet | - | 1:500 | Biolegend | 423113 |  |
| CD4 | OKT4 | FITC | 1:300 | Biolegend | 317407 | AB_571950 |
| CD8 | RPA-T8 | BV510 | 1:300 | Biolegend | 301048 | AB_2561942 |
| TNFa | MAb11 | APC | 1:500 | Biolegend | 502912 | AB_315264 |
| IL-2 | MQ1-17H12 | PE | 1:500 | Biolegend | 500306 | AB_315093 |
| IFNγ | B27 | BB700 | 1:500 | BD Biosciences | 566395 | AB_2744484 |
| 4-1BB Activation Staining |  |  |  |  |  |  |
| Marker | Clone | Fluorophore | Dilution | Supplier | Catalog Nr. | RR_ID |
| Live/Dead | Zombie Violet | - | 1:500 | Biolegend | 423113 |  |
| CD4 | OKT4 | FITC | 1:300 | Biolegend | 317407 | AB_571950 |
| CD8 | RPA-T8 | BV510 | 1:300 | Biolegend | 301048 | AB_2561942 |
| CD137 (4-1BB) | 4B4-1 | AF700 | 1:200 | Biolegend | 309816 | AB_2207742 |
| B7H3 Tumor Cell Staining |  |  |  |  |  |  |
| Marker | Clone | Fluorophore | Dilution | Supplier | Catalog Nr. | RR_ID |
| Live/Dead | DAPI | - | 1.25 ug/ml | ThermoFisher | D1306 |  |
| B7H3 (CD276) | B7RP2 | PE | 1:200 | Biolegend | 351003 | AB_10719959 |
| or |  |  |  |  |  |  |
| Live/Dead | DAPI | - | 1.25 ug/ml | ThermoFisher | D1306 |  |
| B7H3 (CD276) | MGA271 | - | 5 ug/ml | Biointron | n/a | n/a |
| Goat anti-mouse | Poly4053 | PE | 1:200 | Biolegend | 405307 | AB_315010 |
| Tumor digest staining |  |  |  |  |  |  |
| Marker | Clone | Fluorophore | Dilution | Supplier | Catalog Nr. | RR_ID |
| Live/Dead | DAPI | - | 1.25 ug/ml | ThermoFisher | D1306 |  |
| CD4 | SK3 | PE | 1:300 | eBioscience | 12-0047-42 | AB_1518750 |
| CD8 | SK1 | FITC | 1:300 | BioLegend | 344704 | AB_1877178 |
| CD45 | HI30 | BV510 | 0.25 | Biolegend | 304036 | AB_2561940 |
| CAR-T Staining |  |  |  |  |  |  |
| Marker | Clone | Fluorophore | Dilution | Supplier | Catalog Nr. | RR_ID |
| Whitlow Linker | KIP-1 Clone 16 | - | 1 ug/ml | Biointron | n/a | n/a |
| Anti-Rabbit IgG | Polyclonal | PE | 1:300 | Invitrogen | P2771MP | AB_2539845 |

